# Livestock depredation frequency is negatively associated with wolf body mass

**DOI:** 10.64898/2026.10.15.759474

**Authors:** C. Aaron Bott, Roblyn Brown, Daniel R. MacNulty

## Abstract

Foraging theory predicts that behavioral decisions are state-dependent, with individuals expected to alter foraging strategies according to physiological condition. Although livestock depredation by carnivores is often attributed to seasonal changes in prey availability and livestock exposure, less attention has been given to predator state. We evaluated whether seasonal variation in wolf body mass was associated with monthly livestock depredation frequency using 17 years of monitoring data from gray wolves (*Canis lupus*) in Oregon, USA. Using 329 body-mass measurements from 274 wolves, we modeled age- and season-related variation in body mass and generated standardized monthly population-level predictions. Predicted body mass declined through spring and summer, reached its lowest values in late summer, and increased into winter. Monthly depredation frequency decreased as predicted population-level body mass increased; a one-standard-deviation increase in predicted mass was associated with a 36.7% decrease in expected monthly depredation frequency (95% confidence interval: 22.7–48.1% decrease). The negative association persisted across sensitivity analyses based on predicted population-level body mass. When season was modeled explicitly, the association remained negative, but uncertainty increased substantially because predicted body mass and season contained limited independent temporal information. Livestock depredations were therefore most frequent during the portion of the annual cycle when wolves were predicted to have their lowest body mass. The observed association is consistent with state-dependent foraging theory, but the available data could not determine whether body mass itself contributed to depredation frequency or whether other seasonal ecological processes drove the association. Our results identify predator state as a potentially important but understudied component of carnivore–livestock interactions. Integrating measures of predator state with established drivers of conflict, including livestock exposure, husbandry, native prey, and landscape conditions, may help determine whether predator state provides additional information for understanding and anticipating carnivore– livestock conflict.

---

Foraging theory predicts that animals do not make foraging decisions solely in response to external environmental conditions, but also according to their internal state (Stephens and Krebs 1986, Ydenberg et al. 2007, Pyke 2019). Energetic state is a particularly important component of internal state because energy acquisition underlies survival, reproduction, growth, and movement (Gittleman 1985, Brodin and Clark 2007). State-dependent foraging theory therefore predicts that individuals experiencing different energetic conditions may adopt different foraging strategies because the fitness consequences of acquiring—or failing to acquire—resources vary with their current state (Stephens and Krebs 1986, Denryter et al. 2020). Individuals experiencing energetic deficits may consequently alter resource selection, expand their diets, or increase their use of resources that would otherwise be less attractive or less frequently exploited (Ydenberg et al. 2007, Hemingway et al. 2020). For predators, such energetic constraints can influence hunting decisions, prey selection, movement patterns, and responses to risk (Carbone et al. 2007, MacNulty et al. 2020, Ritwika et al. 2024). Although state-dependent foraging is a central concept in behavioral ecology, it has received comparatively little attention in studies of human–wildlife conflict (Boutin 2018).

Direct measurement of physiological state in free-ranging animals remains challenging, particularly over broad spatial and temporal scales, because many physiological variables require invasive instrumentation or technologies poorly suited to long-term deployment in wild animals (Williams et al. 2021). Consequently, studies of free-ranging animals often rely on integrative measures that capture biologically meaningful variation in physiological condition. Body mass is widely measured in free-ranging mammals and integrates variation in structural size, lean tissue, hydration, and energetic reserves (Peig and Green 2010). Although body mass does not directly quantify body composition or identify the physiological mechanisms underlying variation among individuals, it can provide information about aspects of physiological state when differences attributable to sex, age, and structural size are appropriately considered (Peig and Green 2010, Wishart et al. 2024). Body mass may therefore provide a useful, albeit imperfect, proxy for aspects of predator state when evaluating relationships between energetic condition and foraging behavior.

Research on livestock depredation by carnivores has traditionally focused on external drivers of conflict, including livestock abundance, husbandry practices, landscape characteristics, prey availability, and carnivore density (Musiani et al. 2005, Moreira-Arce et al. 2018, Ugarte et al. 2019). These studies have provided valuable insights into the environmental and management conditions associated with the risk of depredation. However, comparatively little attention has been given to how variation in predator state may influence foraging decisions and thereby contribute to depredation risk. Consequently, incorporating predator state may provide additional insight into factors contributing to livestock depredation.

Gray wolves (*Canis lupus*) may provide a useful system for examining relationships between predator state and livestock depredation. Wolf–livestock conflict remains a significant conservation, economic, and management challenge throughout North America and Europe (Musiani et al. 2005, Moreira-Arce et al. 2018, Kiffner et al. 2022). Livestock depredation by wolves also exhibits recurring seasonal patterns, creating an opportunity to examine whether variation in predator state corresponds with seasonal variation in conflict (Musiani et al. 2005, DeCesare et al. 2018, Kiffner et al. 2022).

Here, we evaluated whether variation in wolf body mass, as a population-level proxy for aspects of predator state, was associated with livestock depredation frequency using 17 years of monitoring data (2009–2025) from wolves in Oregon, USA. By evaluating predator state as a potential correlate of livestock depredation, we sought to broaden understanding of carnivore– livestock conflict beyond the external ecological and management factors that have traditionally received much attention. We modeled age-related and seasonal variation in body mass and generated standardized population-level monthly predictions for comparison with monthly livestock depredation frequency. Under state-dependent foraging theory, individuals experiencing poorer energetic conditions may alter foraging behavior as the fitness consequences of acquiring resources change. Although state-dependent foraging theory operates at the level of individuals, our available data allowed us to evaluate the corresponding population-level seasonal pattern. We therefore predicted that livestock depredations would occur more frequently during periods when predicted population-level wolf body mass was lower. Our objective was not to determine whether the body mass of individual wolves predicted their participation in depredation events, but rather whether seasonal variation in predicted population-level body mass corresponded with seasonal variation in livestock depredation frequency.

## Study Area

Our study encompassed Oregon, USA, which includes diverse landscapes ranging from high deserts and shrublands to temperate coniferous forests and mountainous terrain. Gray wolves naturally recolonized Oregon in the early 2000s following population expansion from Idaho and the Northern Rocky Mountains (Oregon Department of Fish and Wildlife [ODFW] 2019). The study covered 17 years (2009–2025), during which wolves were monitored across the state. Livestock within the study system primarily consisted of cattle (*Bos taurus*) and sheep (*Ovis aries*).

## Methods

### Wolf monitoring and data collection

ODFW monitored wolves statewide using Global Positioning System (GPS) and Very High Frequency (VHF) telemetry, aerial tracking, and ground-based surveys. Wolves were captured using foothold traps in spring, summer, and fall, or by aerial darting during winter, and processed under standardized handling protocols (ODFW 2019). Wolf capture and handling were conducted by ODFW as part of its standard wildlife management and monitoring activities. Freshly killed wolves were included if carcasses were intact and weighed immediately after death, thereby minimizing postmortem changes that could affect body mass measurements. Individuals were weighed using calibrated spring or digital scales, and sex and age were determined based on dentition, reproductive morphology, and body size (Kreeger et al. 2003, Mech 2006).

Trained wildlife biologists investigated depredations following reports from livestock producers. Confirmation followed established forensic criteria, including evidence of premortem hemorrhage, characteristic bite marks and tooth spacing, subcutaneous tissue damage, predator sign at the scene, and necropsy indicators distinguishing predation from scavenging (ODFW 2019). We included only incidents confidently classified as confirmed depredations in the analysis.

### Wolf body mass model

We modeled variation in wolf body mass using a generalized additive model (GAM) with a Gaussian error distribution to accommodate nonlinear relationships between body mass, age, and seasonal timing. Body mass (kg) was modeled as a function of sex, capture method, sex-specific nonlinear age trajectories, seasonal timing, and individual identity. Sex-specific growth trajectories were represented using a factor-smooth interaction between age and sex, seasonal variation was represented using a cyclic cubic regression spline for day of year, and wolf identity was included as a random-effect smooth to account for repeated measurements of individuals. Capture method was included to account for systematic differences associated with how wolves entered the body-mass dataset. Wolves of all observed ages were retained because age-related variation in body mass was modeled explicitly rather than controlled by restricting the analysis to a particular age class. This allowed the seasonal component of the model to be estimated after accounting for nonlinear, sex-specific changes in body mass with age.

We evaluated competing model structures using maximum likelihood (ML) and Akaike’s Information Criterion (AIC). Candidate models evaluated support for seasonal variation, capture method, an age × season interaction, and random effects for individual identity, year, and pack. Individual identity was retained to account for repeated measurements of the same wolves. Pack identity provided little additional explanatory information and was not retained in the final model. Similarly, adding year did not improve support for the primary all-wolf model and was therefore excluded from its final structure. The selected model was subsequently refitted using restricted maximum likelihood (REML) for parameter estimation.

To evaluate whether measurements obtained following mortality influenced estimates of seasonal body mass, we repeated the analysis after excluding all postmortem measurements. We compared seasonal predictions from this live-only model with those from the full dataset. Because interannual variation in body mass could provide information for distinguishing body mass from season in subsequent depredation analyses, we also fit versions of the all-wolf and live-only models containing a year random effect and retained these models as sensitivity analyses even when year was not supported in the primary model.

### Population-level body mass prediction

We used the fitted body-mass models to generate population-level predictions across the annual cycle. Predictions were evaluated at the median observed age (1.64 years) and standardized to aerial capture, generated separately for females and males, and then averaged equally across sexes. Holding age constant ensured that seasonal changes in predicted mass reflected the modeled seasonal trajectory rather than seasonal differences in the ages of wolves represented in the sample. Similarly, generating sex-specific predictions before averaging them equally prevented temporal variation in the sampled sex ratio from determining the population-level trajectory. Thus, the resulting predictor represented seasonal variation in expected body mass for a standardized wolf population rather than the raw mean mass of wolves sampled in a given month. Individual-wolf random effects were excluded from prediction so that the resulting trajectory represented the expected population-level seasonal pattern rather than the expected mass of particular sampled individuals. Daily predictions were aggregated to monthly means and standardized (mean = 0, SD = 1) before inclusion in depredation models.

Because year was not included in the primary body-mass model, its predicted seasonal trajectory was common among years. We therefore generated a second set of predictions from models containing a year random effect. For these sensitivity predictions, individual effects were excluded while estimated year effects were retained, allowing monthly predicted body mass to vary among years.

### Relationship between body mass and livestock depredation

Confirmed livestock depredations were aggregated by year and month from 2009–2025, including months in which no confirmed depredations were recorded. We first evaluated the association between predicted population-level body mass and monthly depredation frequency using a negative binomial generalized linear mixed model (GLMM), with standardized predicted body mass as the explanatory variable and random intercepts for year and month. The negative binomial distribution was used to accommodate overdispersion in depredation counts and was better supported than a Poisson model. This analysis tested whether livestock depredations occurred more frequently during periods when wolves were predicted to have lower body mass. Because both the body-mass predictor and depredation response were evaluated at the population-month scale, this analysis tested a temporal population-level association rather than whether the body mass of an individual wolf predicted its probability of participating in a depredation. Individual participation in confirmed depredations was generally unknown, and depredations may involve multiple members of a pack; consequently, individual-level body mass could not be directly linked to individual foraging decisions.

Because predicted body mass itself exhibited a pronounced annual cycle, the primary GLMM could not by itself distinguish an association with body mass from other ecological processes that also vary seasonally. We therefore fit a more conservative negative binomial generalized additive mixed model (GAMM) containing standardized predicted body mass, a cyclic smooth for month, and a random effect for year. This model tested whether predicted body mass explained variation in depredation frequency beyond the broad seasonal pattern in depredations. We evaluated dependence between the body-mass predictor and seasonal smooth using concurvity diagnostics and compared the magnitude and uncertainty of the body-mass coefficient between the GLMM and season-adjusted GAMM.

We repeated both depredation models using body-mass predictions derived exclusively from live wolves and using year-specific body-mass predictions. These analyses evaluated whether inclusion of postmortem measurements or the limited interannual variation in body mass affected inference. To propagate uncertainty associated with estimated body mass, we additionally conducted 1,000 Monte Carlo simulations in which coefficients were sampled from the multivariate sampling distribution of the fitted body-mass model, new population-level body-mass predictors were generated, and the corresponding depredation model was refitted.

#### Additional Analyses of Body Mass

We conducted an additional analysis intended to isolate variation in body mass that was not attributable to the expected seasonal cycle. For each measured wolf, we calculated residual body mass as the difference between observed and model-predicted mass. Positive residuals therefore represented wolves that were heavier than expected given the modeled predictors, whereas negative residuals represented wolves that were lighter than expected. Residual body mass was summarized at the pack × year level and related to annual depredation frequency using negative binomial models containing pack and year random effects. Because body-mass sampling within pack-years was sparse, we evaluated datasets requiring at least one, two, or three mass observations per pack-year to determine whether inference was sensitive to within-pack sampling intensity.

We also evaluated whether finer spatial matching between measured wolves and depredation events at the pack level provided additional information. However, temporal overlap between body-mass measurements and depredations within individual packs was limited, and pack identity explained little additional variation in body mass. These analyses were therefore considered exploratory and were not used as the primary basis for inference.

Finally, because capture method explained variation in measured body mass, we tested whether wolves lethally removed by management following livestock conflicts differed systematically in body mass from other sampled wolves. Management removals were identified from mortality records associated with trapping, ground shooting, or aerial removal; wolves recovered dead from other causes were excluded from this classification. Because management removal did not establish that an individual participated in a particular depredation event, this analysis was interpreted only as an exploratory comparison.

We considered incorporating an exposure offset representing wolves’ opportunity to encounter livestock, but the available data did not contain a consistent measure of livestock availability across pack territories and months. Numbers of livestock recorded during individual depredation investigations represented animals associated with specific investigations rather than livestock exposure to wolves and were therefore not used as an offset. Model adequacy for negative binomial depredation models was evaluated using simulation-based residual diagnostics, including tests for residual uniformity, dispersion, and zero inflation.

## Results

### Wolf body mass

We recorded 329 body-mass measurements from 274 individual wolves (Fig. 1). Of these, 228 wolves were measured once, 37 twice, and 9 three times. The dataset contained 184 initial handling records, 30 live recapture records, and 115 measurements obtained following mortality. The 329 measurements included 163 female and 166 male handling events. The live-only sensitivity dataset contained 214 measurements from 187 individual wolves.

**Figure 1.**
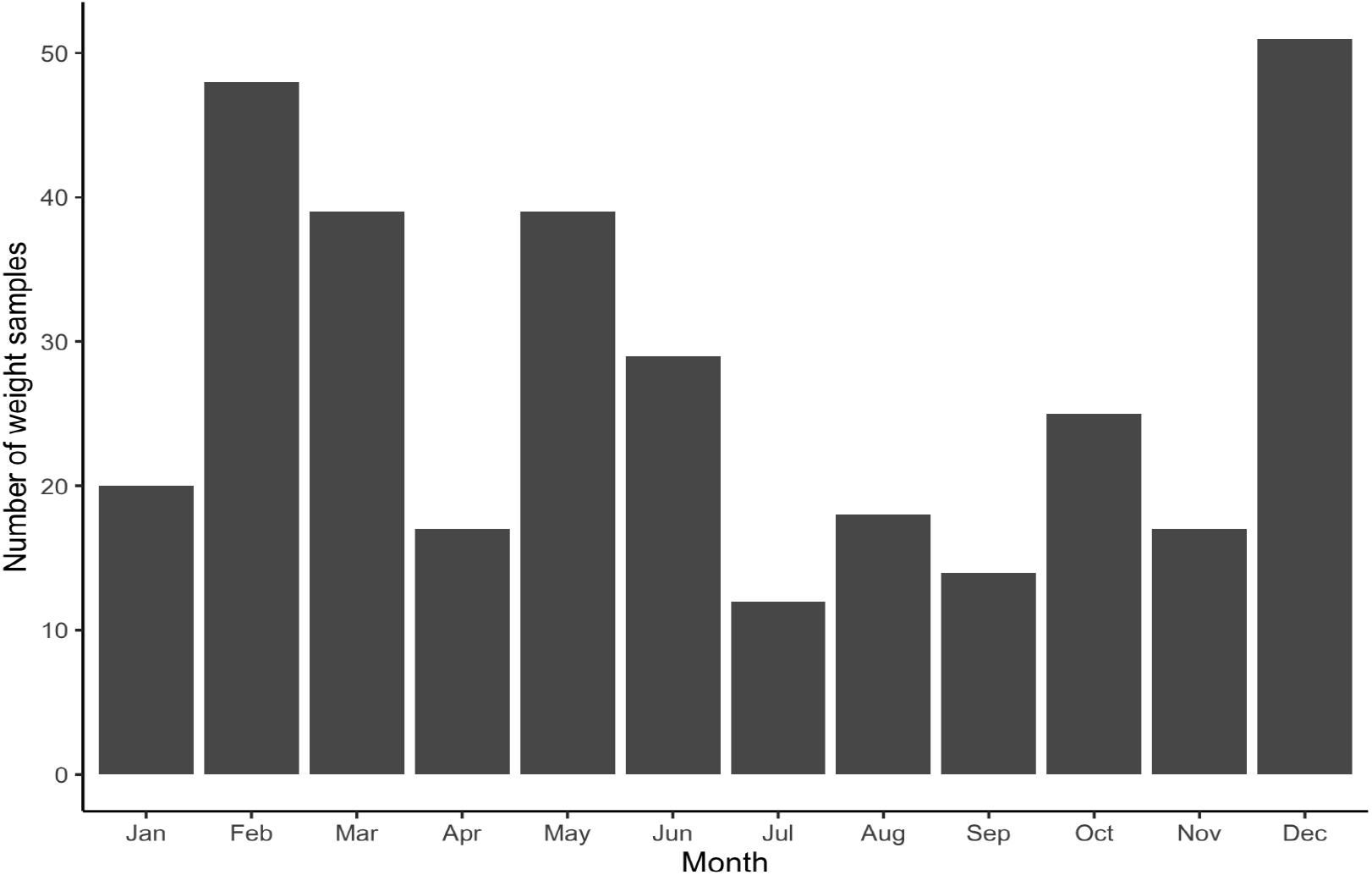
Monthly distribution of wolf body-mass measurements used to model age-related and seasonal variation in body mass in Oregon gray wolves, 2009–2025. A total of 329 body-mass measurements were collected from 274 individual wolves, including 46 individuals that were measured more than once (37 measured twice and 9 measured three times).

The best-supported body-mass model included sex, capture method, sex-specific nonlinear age trajectories, seasonal variation, and individual wolf identity (Table 1). Seasonal variation and capture method were strongly supported: removing season increased AIC by 70.38, whereas removing capture method increased AIC by 11.37. An age × season interaction received little support (ΔAIC = 59.91). Adding year to the primary all-wolf model did not improve model support relative to the more parsimonious model without year (ΔAIC = 1.64 relative to the model without year), and pack identity similarly provided little additional explanatory information. Year and pack were therefore excluded from the primary model, although year was retained in subsequent sensitivity analyses.

**Table 1.**
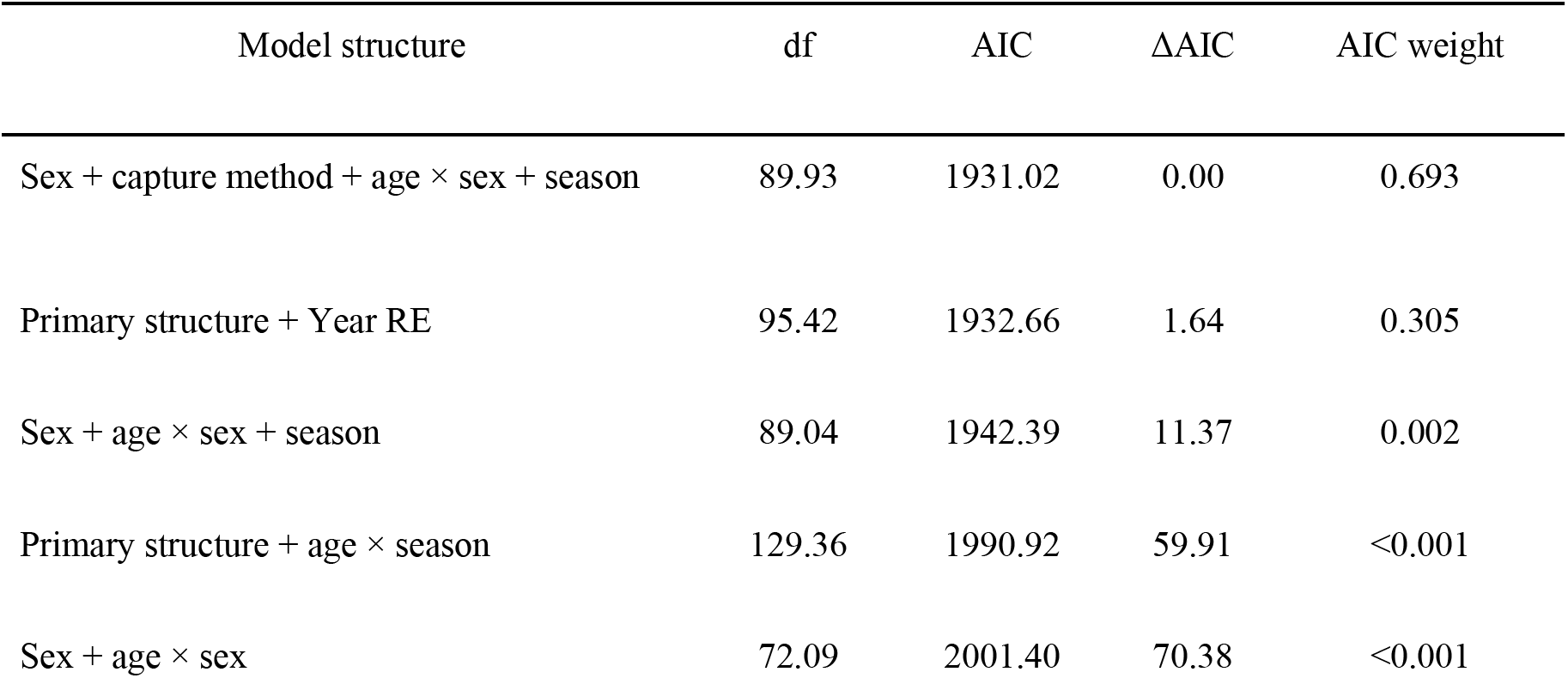
Model selection results for Gaussian generalized additive models describing variation in gray wolf body mass in Oregon, USA, 2009–2025. Models were fit using maximum likelihood (ML) and compared using Akaike’s Information Criterion (AIC). Individual wolf identity was included as a random-effect smooth in all models shown to account for repeated measurements of the same wolves. The primary model included sex, capture method, sex-specific nonlinear age trajectories, seasonal variation, and individual identity. A model additionally containing a year random effect was retained as a sensitivity analysis because it received some support but did not improve AIC relative to the more parsimonious primary structure.

The final body-mass model explained 81.5% of deviance in wolf body mass (Table 2). Males were approximately 8.0 kg heavier than females after accounting for other model terms, and growth trajectories were strongly nonlinear and differed between sexes (Fig. 2). Body mass exhibited a pronounced seasonal cycle (edf = 2.47, *P* = 0.002), declining through spring and summer to its lowest predicted values in late summer before increasing into winter. Because age and sex were accounted for in the body-mass model and held constant or standardized when generating population-level predictions, the predicted seasonal trajectory was not attributable to seasonal changes in the age or sex composition of sampled wolves.

**Figure 2.**
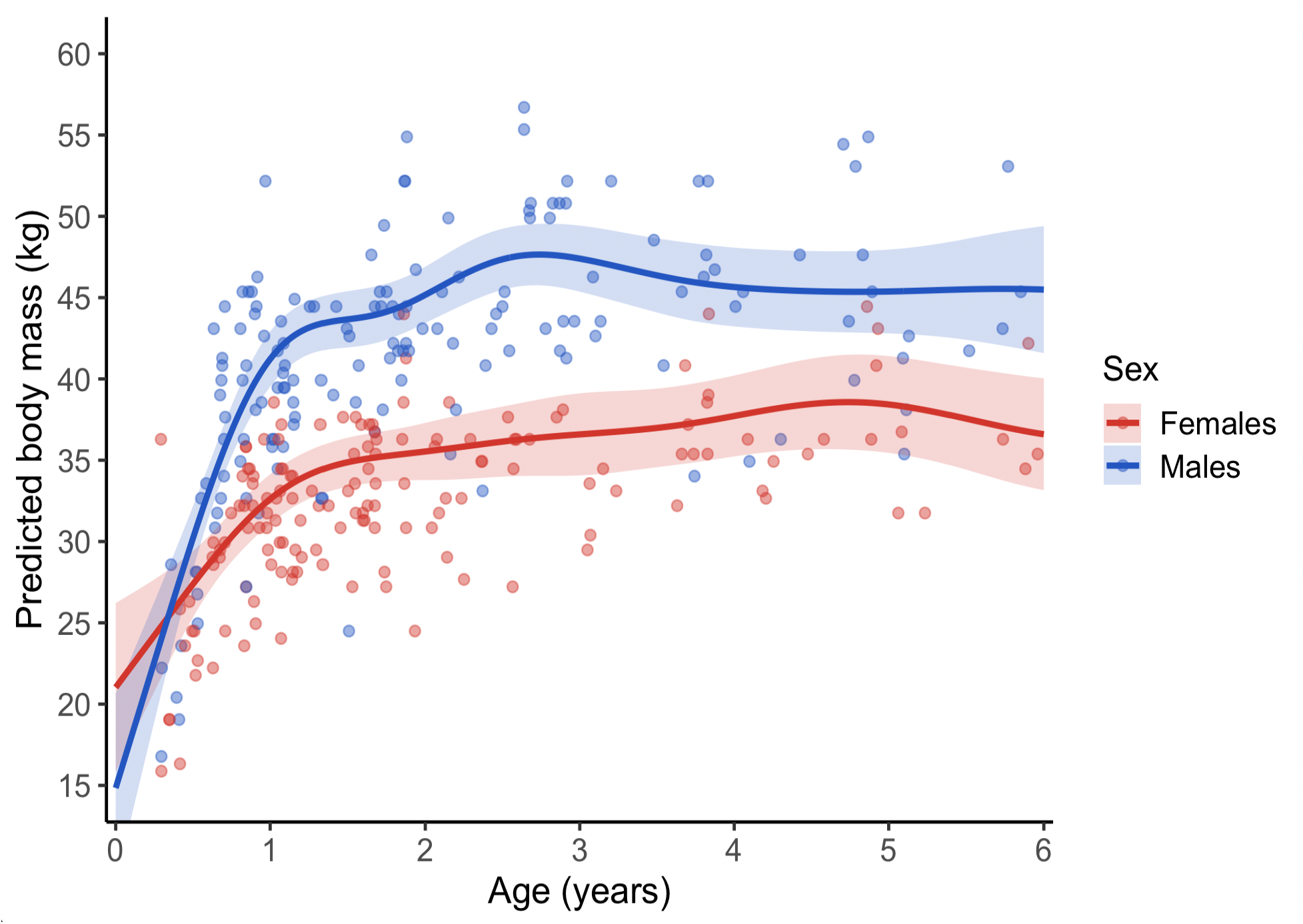
Predicted body mass (kg) as a function of age (years) for female and male gray wolves in Oregon, USA, 2009–2025, derived from the final Gaussian generalized additive mixed model. Points represent observed body-mass measurements from individual wolves. Solid lines represent model-predicted mean body mass as a function of age for each sex, standardized to aerial capture, with seasonal and individual-wolf effects centered or excluded; shaded ribbons represent 95% confidence intervals around the predicted mean. Thus, points represent individual measurements, whereas lines represent population-level model predictions rather than trajectories of individual wolves. The plot is restricted to wolves ≤6 years old because observations at older ages were sparse; all observations were retained in model fitting and subsequent analyses.

**Table 2.**
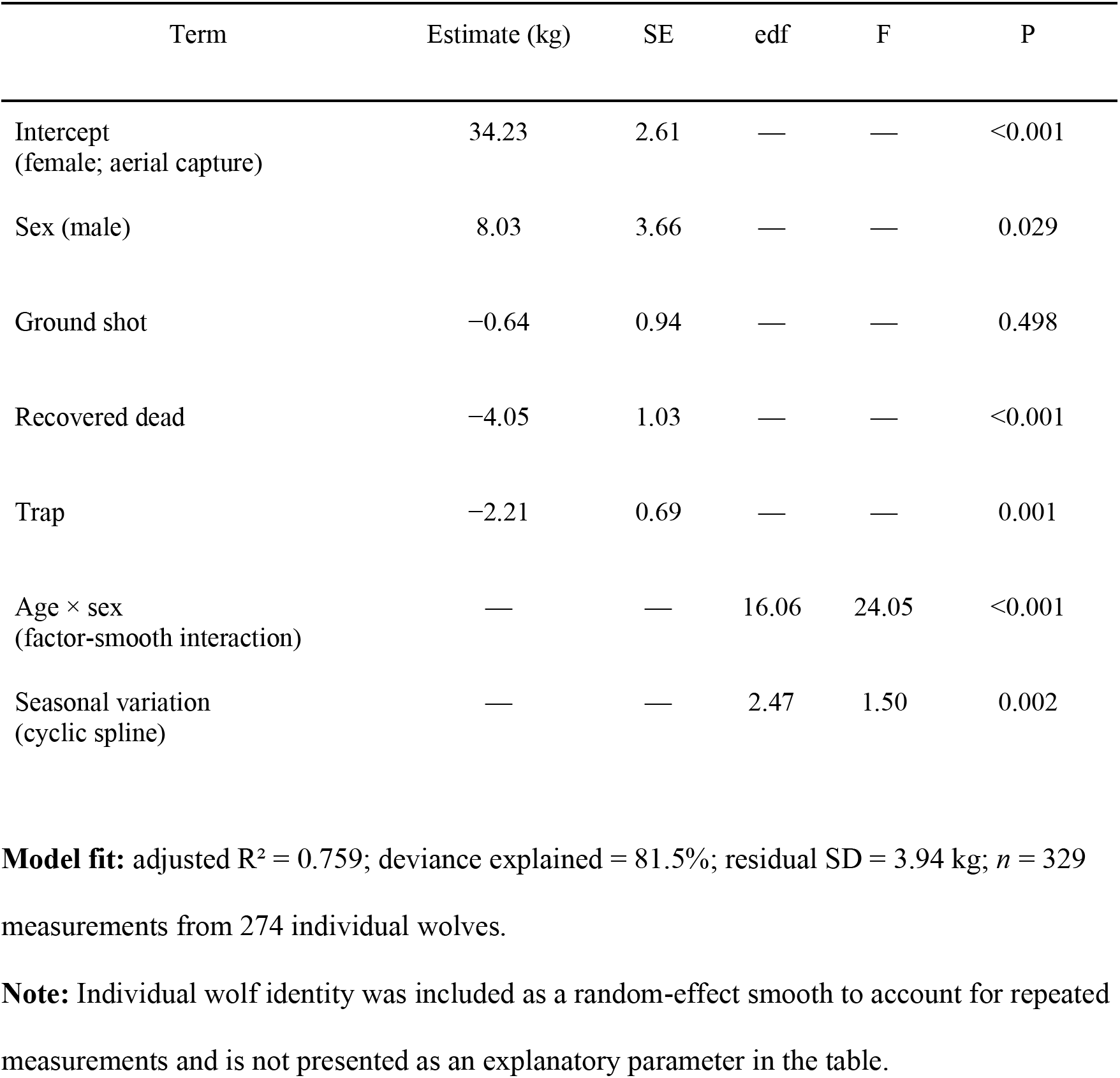
Parameter estimates and smooth-term summaries from the final Gaussian generalized additive mixed model describing gray wolf body mass in Oregon, USA, 2009–2025. The model included sex, capture method, sex-specific nonlinear age trajectories, seasonal variation modeled with a cyclic spline of day of year, and individual wolf identity as a random effect to account for repeated measurements. Aerial capture was the reference category for capture method, and females were the reference sex. The final model was fitted using restricted maximum likelihood.

Capture method explained additional variation in body mass. Relative to aerial-captured wolves, wolves classified as recovered dead were approximately 4.05 kg lighter (SE = 1.03 kg, P < 0.001), and trap-captured wolves were approximately 2.21 kg lighter (SE = 0.69 kg, P = 0.001), after accounting for age, sex, seasonal timing, and repeated measurements. The estimated difference between ground-shot (i.e., chemically immobilized or lethally shot from the ground using a firearm) wolves and aerial-captured wolves was small relative to its uncertainty.

The live-only model explained 81.4% of the deviance in wolf body mass and produced a similar seasonal pattern (Fig. S1), indicating that mortality-derived measurements did not drive the estimated seasonal cycle.

When year was added to the all-wolf model, interannual variation in body mass was modest (year random-effect SD = 0.59 kg compared with individual-wolf SD = 1.87 kg), increasing explained deviance from 81.5% to 82.1%. Nevertheless, we retained these year-specific predictions to determine whether even modest interannual variation altered inference about depredation frequency.

### Relationship between body mass and livestock depredation

We recorded 548 confirmed livestock depredation events across 204 monthly observations from 2009–2025; 70 months contained no confirmed depredations.

Monthly livestock depredation frequency exhibited a pronounced seasonal pattern that corresponded inversely with predicted population-level body mass (Fig. 3). In the primary negative binomial GLMM, monthly livestock depredation frequency was negatively associated with predicted population-level body mass (Fig. 4; Table 3) (β = −0.457, SE = 0.102; IRR = 0.633, 95% CI = 0.519–0.773), indicating that a 1-SD increase in predicted body mass was associated with a 36.7% decrease in expected monthly depredation frequency. Thus, depredations occurred more frequently during periods of the annual cycle when standardized population-level wolf body mass was predicted to be lower.

**Figure 3.**
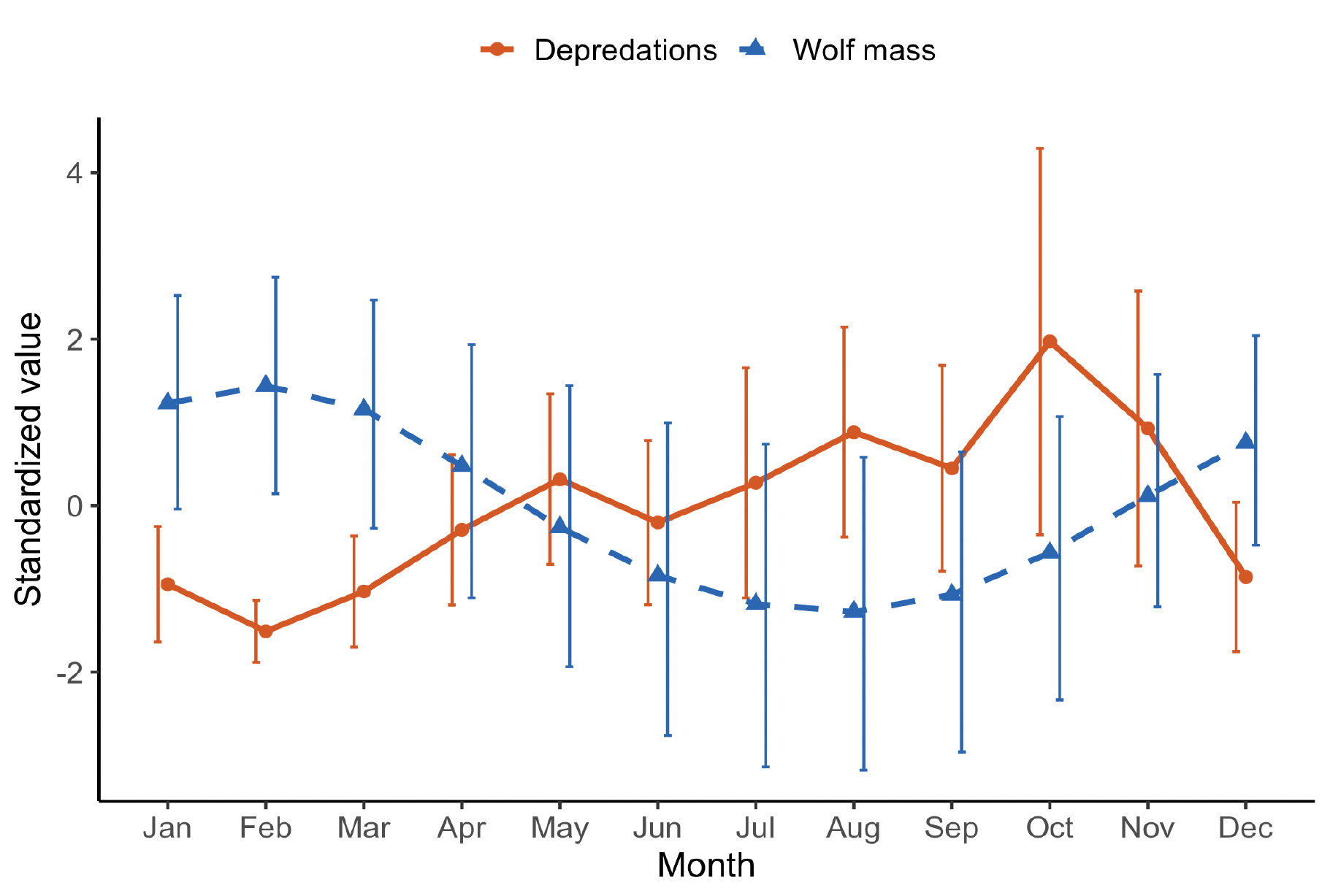
Monthly correspondence between predicted population-level wolf body mass and livestock depredation frequency in Oregon, USA, 2009–2025. The blue dashed line with triangular points represents standardized monthly mean predicted body mass from the final Gaussian generalized additive mixed model, with age held constant at the median observed value (1.64 years), capture method standardized to aerial capture, sexes averaged equally, and individual-wolf effects excluded. Because year was not included in the final body-mass model, predicted mass represents the common seasonal trajectory across years. Error bars for body mass represent model-based uncertainty propagated using 1,000 coefficient draws. The orange solid line with circular points represents mean monthly confirmed livestock depredation frequency across years, with error bars representing among-year variation (±1.96 SE).

**Figure 4.**
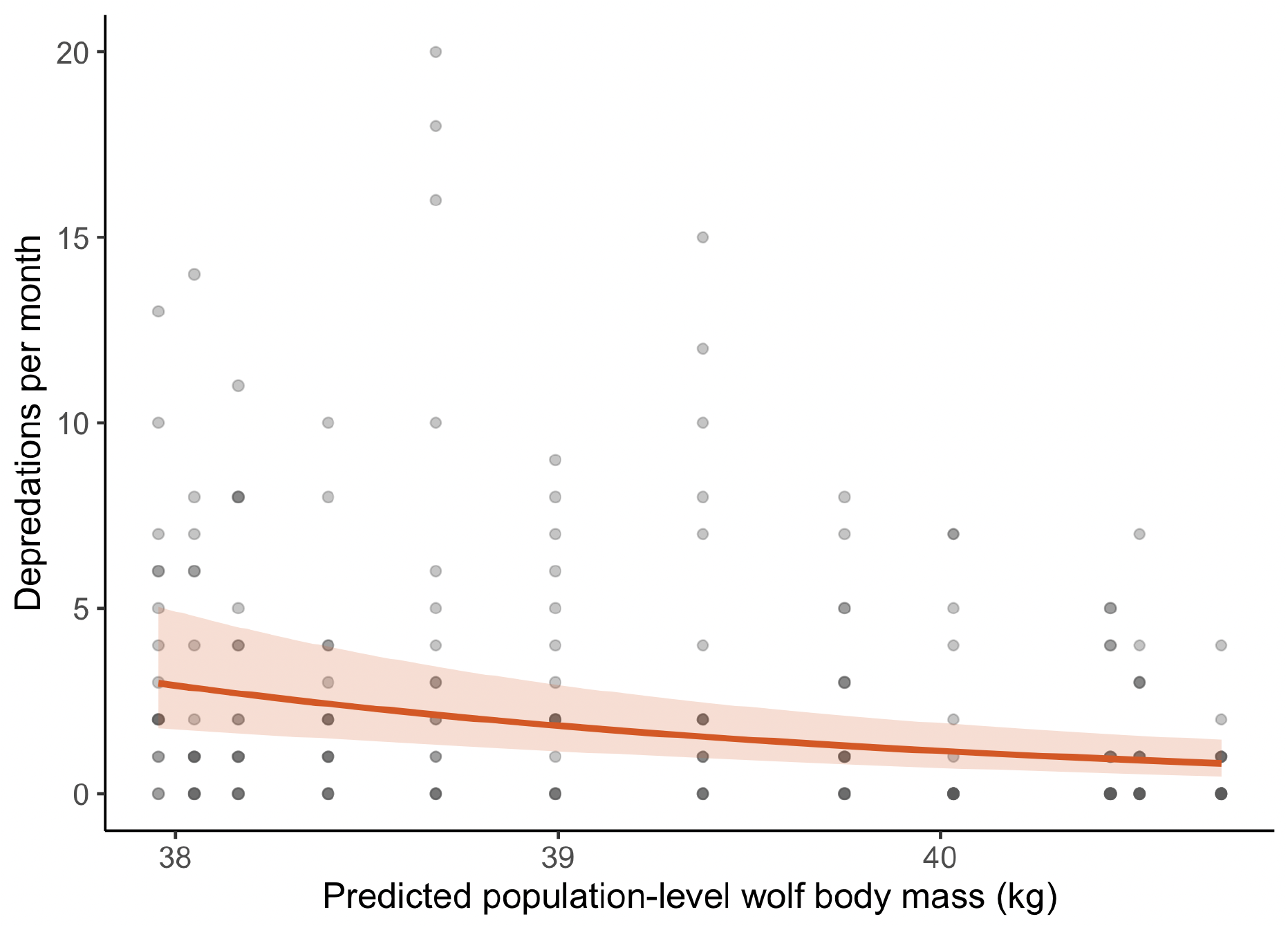
Relationship between standardized predicted population-level wolf body mass and confirmed monthly livestock depredation frequency in Oregon, USA, 2009–2025. Predicted body mass represents monthly mean predictions from the final Gaussian generalized additive mixed model, evaluated at the median observed age (1.64 years), standardized to aerial capture, averaged equally across sexes, and excluding individual-wolf effects. Because year was not included in the body-mass model used for these predictions, predicted mass represents a common seasonal trajectory across years. The solid line represents predictions from the negative binomial generalized linear mixed model relating standardized predicted body mass to monthly depredation frequency, with random intercepts for year and month; the shaded band represents the 95% confidence interval. Points represent observed depredation counts for the 204 year-month observations.

**Table 3.** Relationship between standardized predicted gray wolf body mass and monthly livestock depredation frequency in Oregon, USA, 2009–2025. Results are shown for the primary negative binomial generalized linear mixed model (GLMM) and the season-adjusted negative binomial generalized additive mixed model (GAMM). The GLMM included random intercepts for year and month. The GAMM included predicted body mass, a cyclic smooth for month, and a random effect for year. Models were repeated using body-mass predictions derived from all measurements and from live wolves only. Incidence rate ratios (IRR) represent the multiplicative change in expected monthly depredation frequency associated with a 1-SD increase in predicted body mass.

**A. Deterministic model estimates**
| Body-mass dataset | Model | $\beta$ | SE | $z$ | $P$ | IRR | 95% CI (IRR) |
| --- | --- | --- | --- | --- | --- | --- | --- |
| All wolves | Primary NB GLMM | −0.457 | 0.102 | −4.49 | <0.001 | 0.633 | 0.519–0.773 |
| Live wolves only | Primary NB GLMM | −0.402 | 0.112 | −3.59 | <0.001 | 0.669 | 0.537–0.833 |
| All wolves | Season-adjusted NB GAMM | −0.591 | 0.430 | −1.37 | 0.170 | 0.554 | 0.238–1.287 |
| Live wolves only | Season-adjusted NB GAMM | −0.269 | 0.531 | −0.51 | 0.612 | 0.764 | 0.270–2.162 |

**B. Monte Carlo uncertainty propagation (n = 1000)**
| Body-mass dataset | Model | Mean $\beta$ | SD | 95% simulation interval ($\beta$) | Mean IRR | 95% simulation interval (IRR) | $\beta < 0$ | $\beta < 0$ and $P < 0.05$ |
| --- | --- | --- | --- | --- | --- | --- | --- | --- |
| All wolves | Primary NB GLMM | -0.417 | 0.073 | -0.500 to -0.243 | 0.661 | 0.606–0.784 | 99.8% | 96.0% |
| Live wolves only | Primary NB GLMM | -0.331 | 0.153 | -0.475 to 0.149 | 0.728 | 0.622–1.161 | 94.8% | 79.6% |
| All wolves | Season-adjusted NB GAMM | -0.280 | 0.260 | -0.672 to 0.252 | 0.783 | 0.511–1.286 | 83.6% | 22.3% |
| Live wolves only | Season-adjusted NB GAMM | -0.149 | 0.281 | -0.635 to 0.349 | 0.896 | 0.530–1.418 | 66.1% | 7.7% |

The association persisted when postmortem measurements were excluded from the body-mass model. Using the live-only mass predictor, the primary GLMM produced a similar negative association (β = −0.402, SE = 0.112; IRR = 0.669, 95% CI = 0.537–0.833). Explicitly accounting for season substantially increased uncertainty in the estimated body-mass effect. In the negative binomial GAMM containing both predicted body mass and a cyclic smooth for month, the body-mass coefficient remained negative and increased in magnitude from β = −0.457 in the primary GLMM to β = −0.591, SE = 0.430 (IRR = 0.554, 95% CI = 0.238–1.287) in the season-adjusted GAMM. However, the confidence interval widened considerably and encompassed a broad range of associations, from a substantial negative association to effects in the opposite direction. The standard error of the body-mass coefficient increased more than fourfold relative to the primary GLMM. The live-only GAMM likewise produced a negative but imprecisely estimated association (β = −0.269, SE = 0.531; IRR = 0.764, 95% CI = 0.270–2.162).

Concurvity diagnostics were consistent with overlap between the seasonal body-mass predictor and the cyclic month term. Maximum (worst) concurvity approached 1.0, although observed and estimated concurvity measures were lower. Together with the pronounced increase in uncertainty surrounding the body-mass coefficient after explicit seasonal adjustment, these results indicated that the available data contained limited independent variation to reliably partition the contributions of predicted body mass and season.

Allowing body mass to vary among years did not resolve this limitation. Using year-specific predictions, the primary GLMM again produced a negative association (β = −0.473, SE = 0.104; IRR = 0.623, 95% CI = 0.508–0.764), equivalent to a 37.7% decrease in expected depredation frequency per 1-SD increase in predicted body mass. The corresponding season-adjusted GAMM likewise produced a negative but less precisely estimated association (β = −0.576, SE = 0.377; IRR = 0.562, 95% CI = 0.269–1.176). Thus, introducing observed interannual variation in predicted body mass did not materially alter the overall pattern.

Monte Carlo simulations propagating uncertainty in predicted body mass produced the same general pattern. Across 1,000 retained simulations, the mean body-mass coefficient in the primary GLMM was −0.417 using all body-mass measurements and −0.331 using live-only measurements. Corresponding Monte Carlo mean coefficients from the season-adjusted GAMM were closer to zero (mean β = −0.280 and −0.149, respectively). Thus, propagating uncertainty in estimated body mass did not materially alter the negative descriptive association observed in the primary models; rather, estimates became substantially less informative when the shared seasonal structure of body mass and depredation was modeled explicitly.

### Additional analyses of body mass

Associations between residual body mass and livestock depredation frequency varied across the sampling thresholds evaluated. Effect estimates were small, varied in direction, and were estimated with substantial uncertainty. Moreover, available replication declined substantially when greater within-pack sampling was required: only 46 pack-years contained at least two body-mass measurements and 25 contained at least three. Finer pack-level analyses were similarly constrained by sparse temporal overlap between measured wolves and depredation events and produced little additional information about the relationship between body mass and depredation.

The estimated difference in body mass between wolves lethally removed by management following livestock conflicts and other sampled wolves was small relative to its uncertainty (β = 0.165 ± 0.653 SE).

Simulation-based residual diagnostics provided no evidence of problematic dispersion, zero inflation, or residual nonuniformity in the negative binomial depredation models. Thus, the loss of precision in the season-adjusted models was most consistent with limited independent variation between body mass and season rather than an obvious failure of model fit.

## Discussion

We found a strong temporal correspondence between seasonal variation in wolf body mass and livestock depredation frequency. Depredations were most frequent during periods when wolves were predicted to have their lowest body mass. However, body mass and depredation frequency both exhibited pronounced seasonal structure. When season was modeled explicitly using a flexible cyclic smooth, the estimated body-mass effect remained negative and increased in magnitude, but its confidence interval widened considerably. Thus, the season-adjusted estimate remained consistent with a substantial negative association while also encompassing little or no association and effects in the opposite direction. Across the primary and sensitivity analyses based on predicted population-level body mass, estimates were directionally consistent, although their precision varied depending on how much independent information was available to distinguish body mass from season. Together, these results demonstrate a consistent population-level temporal association between depredation frequency and predicted wolf body mass, while leaving unresolved whether body mass itself contributes to that association.

This distinction matters because the observed pattern is nevertheless consistent with state-dependent foraging theory. State-dependent models predict that foraging decisions depend not only on resource availability but also on the forager’s physiological state (Stephens and Krebs 1986, Ydenberg et al. 2007, Denryter et al. 2020). As energetic reserves decline, the fitness consequences of failing to acquire food may increase, potentially altering an animal’s willingness to accept costs or risks associated with particular foraging opportunities (Blecha et al. 2018). Individuals in poorer physiological condition may therefore value food differently than individuals in better condition and alter resource use accordingly (Galicia et al. 2020). Under this hypothesis, energetic state could influence the circumstances under which wolves use livestock as prey, particularly when the availability or profitability of native prey changes relative to domestic prey (Morehouse and Boyce 2011, Nelson et al. 2016). This hypothesis does not compete with established external determinants of carnivore–livestock conflict, including native prey availability, livestock husbandry, grazing regimes, landscape characteristics, and opportunities for predator–livestock encounters (Musiani et al. 2005, Moreira-Arce et al. 2018, Ugarte et al. 2019, Kiffner et al. 2022). Rather, physiological state could modify how predators respond to those environmental opportunities. The temporal correspondence observed here is consistent with this hypothesis, but does not establish the underlying behavioral mechanism.

The scale of inference in our analysis is population-level rather than individual. Both predicted body mass and depredation frequency were evaluated at the population-month scale, whereas state-dependent foraging decisions occur at the level of individuals or social groups. Wolves hunt and defend territories cooperatively, and livestock depredations may involve one or several pack members. Body mass, by contrast, was measured from individual wolves at discrete points in time. Population-level predictions allowed us to characterize the broad seasonal body-mass pattern in Oregon wolves, but they cannot establish the physiological condition of the particular animals involved in a depredation. Consequently, our results do not demonstrate that an individual wolf with lower body mass is more likely to depredate livestock than a heavier wolf. Rather, they show that depredations were more frequent during periods when standardized population-level body mass was predicted to be lower.

Body mass provides an imperfect but biologically informative proxy for aspects of physiological state. Body mass combines structural size, fat, and lean tissue and therefore cannot be interpreted as a direct measure of energetic reserves. Moreover, comparisons of mass among animals of different ages or structural sizes can confound variation in energetic state with normal growth and scaling (Peig and Green 2010). We addressed these sources of variation by explicitly modeling sex and nonlinear age-related growth before estimating seasonal changes in body mass. Recent work in wild mammals further suggests that this approach is not necessarily inferior to commonly used morphological body-condition indices: Wishart et al. (2024) found that such indices were no better than body mass alone for predicting fat or lean mass across the species and ecological contexts they examined. We therefore interpret modeled seasonal variation in body mass as a population-level proxy for variation potentially associated with physiological state rather than a direct measure of energetic reserves. The seasonal pattern itself was pronounced: predicted wolf body mass declined from winter through spring and summer, reached its lowest values in late summer, and subsequently increased into winter. This finding is consistent with other North American and European studies (Seal and Mech 1983, Mech and Buhl 2020, Strömgren 2023). Numerous processes could contribute to this cycle. Seasonal changes in native prey vulnerability and profitability, reproductive investment, pup provisioning, movement and territorial costs, environmental conditions, and prey acquisition could all affect wolf energy balance (Kreeger et al. 2003, Gable et al. 2018).

The pronounced seasonality of both body mass and livestock depredation creates the principal challenge for interpreting their association because other seasonal processes could contribute to both patterns. Seasonal changes in livestock distribution and vulnerability, grazing practices, wolf reproductive behavior, native prey vulnerability, or encounters between wolves and livestock could increase depredation during the same period that wolf mass declines. Because standardized monthly livestock exposure within occupied wolf territories was unavailable, we could not directly distinguish changing opportunity from changing predator state. We therefore evaluated the association using multiple complementary analyses designed to test its sensitivity to alternative sources of variation and uncertainty. Across sensitivity analyses based on predicted population-level body mass—including exclusion of postmortem measurements, propagation of prediction uncertainty, and incorporation of interannual variation—the estimated association remained negative. Explicitly modeling season likewise did not attenuate the point estimate; instead, its uncertainty expanded substantially because predicted body mass and season contained much of the same temporal information. Additional analyses using pack-level matching and residual body mass were constrained by sparse replication and provided limited independent information. Thus, the available data contained insufficient independent variation to determine whether body mass itself contributed to depredation frequency or whether the observed association reflected other seasonal processes.

Resolving these alternatives will require data collected at a scale capable of directly linking predator state with subsequent foraging behavior. Denser repeated measurements—or alternative measures of condition—at the level of individual wolves or packs could provide variation in physiological state that can be evaluated independently of the average seasonal trajectory. Integrating such measurements with GPS movement data, kill-site investigations, native prey dynamics, and spatially and temporally explicit measures of livestock exposure could test whether wolves or packs in poorer-than-expected condition subsequently alter prey selection or depredation risk. Such data would provide a more direct test of state-dependent foraging than is possible from population-level seasonal correspondence alone.

The population-level nature of this association also defines its current management relevance. The seasonal association documented here should not be interpreted as evidence that nutritionally stressed wolves are responsible for livestock depredations or that improving wolf energetic condition would reduce conflict. Nor does it displace the established importance of livestock husbandry, native prey, landscape context, and opportunities for wolf–livestock encounters (Ugarte et al. 2019). Across 17 years, livestock depredations in Oregon were most frequent during the portion of the annual cycle when wolves were predicted to have their lowest body mass, and the negative association was consistent across the primary and sensitivity analyses based on predicted population-level body mass. However, the shared seasonality of body mass and depredation prevents attributing this pattern to energetic state alone.

## Research Implications

The negative seasonal association between predicted wolf body mass and livestock depredation frequency suggests that predator state warrants greater consideration in studies of carnivore–livestock conflict. Although shared seasonality prevents us from determining whether energetic state independently contributes to depredation, our findings highlight the importance of evaluating physiological condition alongside established ecological and management drivers. Future research integrating repeated measurements of individual or pack condition with subsequent foraging behavior, livestock exposure, native prey availability, and husbandry practices could help determine whether predator state provides additional explanatory or predictive information about livestock depredation.

## Acknowledgements

We thank ODFW for long-term wolf monitoring and for supporting the collection of field data used in this study. We are especially grateful to the many current and former ODFW biologists and field staff who contributed to wolf monitoring and data collection during the study period. Additionally, we thank the field staff of the U.S. Fish and Wildlife Service, the Confederated Tribes of Warm Springs, and the Confederated Tribes of the Umatilla Indian Reservation for their contributions to wolf monitoring and data collection.

## Author Contributions

C. Aaron Bott designed the study, collected data, analyzed data, and wrote the paper; Daniel R. MacNulty helped design the study, contributed to data analysis, and edited the paper; Roblyn Brown supervised fieldwork and oversaw data collection during the study period.

## Conflict of Interest

The authors declare no conflicts of interest.

## Ethics Statement

Wolf capture and handling were conducted by ODFW as part of its statutory wildlife management and monitoring activities. These agency activities did not require separate field research permits or institutional animal care and use committee (IACUC) approval. Wolves were handled according to protocols approved by ODFW wildlife veterinarians and recommendations of the American Society of Mammalogists (Sikes et al. 2016).

## Statement of Inclusion

This study was conducted within Oregon, USA, and includes authors affiliated with institutions actively involved in wolf management within the study region. The research design and long-term monitoring program were developed in collaboration with state and federal wildlife agencies responsible for wolf conservation. Field data were collected through coordinated efforts among agency biologists, Tribal authorities, and local wildlife professionals. Relevant literature produced by researchers working within the region was cited where appropriate. Study findings will be shared with state and federal managers to inform ongoing conservation and management efforts.

## Funding

This research received no external funding.

## Data Availability

Upon acceptance, the data and R code necessary to reproduce the analyses will be available in the Dryad Digital Repository. Certain identifying and spatial information will be excluded from the publicly archived datasets to protect sensitive information.

## Notes

### Competing Interest Statement

The authors have declared no competing interest.

