## Supplemental Figure 1, Supplemental Table 1 for "Livestock depredation frequency is negatively associated with wolf body mass"

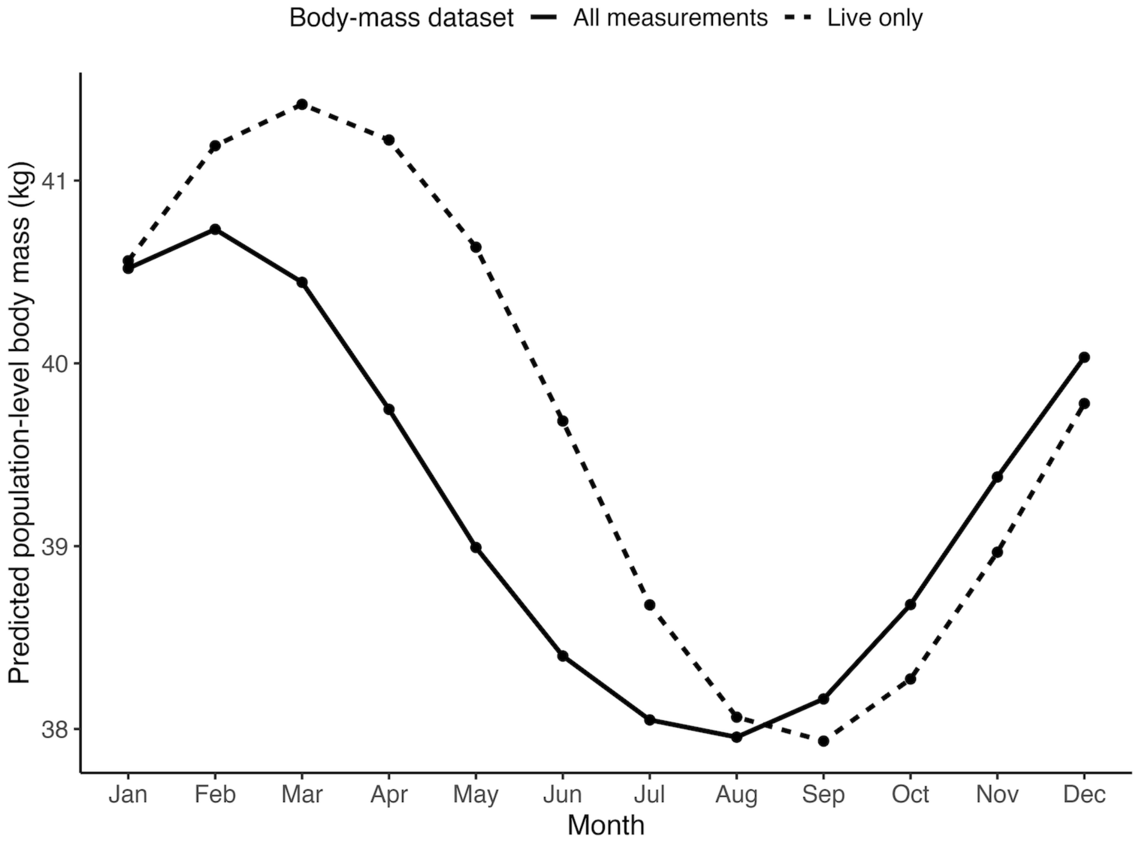


**Figure S1.** Sensitivity analysis comparing predicted seasonal variation in population-level wolf body mass using all body-mass measurements and live-wolf measurements only in Oregon, USA, 2009–2025. Lines represent monthly mean predicted body mass evaluated at the median observed age (1.64 years), standardized to aerial capture, averaged equally across sexes, and excluding individual-wolf effects. The solid line represents predictions from the body-mass model using all 329 measurements, whereas the dashed line represents predictions from a model refit after excluding 115 postmortem measurements (n = 214 live-wolf measurements). Both models produced a similar seasonal pattern, with body mass declining through spring and summer, reaching its lowest values in late summer to early fall, and increasing through fall and winter.

**Table S1.** Sensitivity analyses evaluating the relationship between standardized predicted gray wolf body mass and monthly livestock depredation frequency in Oregon, USA, 2009–2025. Year-specific body-mass predictions were generated from body-mass models containing a year random effect, thereby allowing predicted mass to vary among years rather than repeating an identical seasonal trajectory annually. Results are shown for the negative binomial generalized linear mixed model (GLMM) and the season-adjusted negative binomial generalized additive mixed model (GAMM). The GLMM included random intercepts for year and month; the GAMM included a cyclic smooth for month and a random effect for year. Models were repeated using all body-mass measurements and live-only measurements. β represents the coefficient for standardized predicted body mass; SE = standard error; IRR = incidence rate ratio; CI = confidence interval.

| Body-mass dataset | Prediction structure | Depredation model | β | SE | *P* | IRR | 95% CI |
|---|---|---|---|---|---|---|---|
| All wolves | Year-specific mass | NB GLMM | −0.473 | 0.104 | <0.001 | 0.623 | 0.508–0.764 |
| All wolves | Year-specific mass | Season-adjusted NB GAMM | −0.576 | 0.377 | 0.126 | 0.562 | 0.269–1.176 |
| Live wolves only | Year-specific mass | NB GLMM | −0.461 | 0.121 | <0.001 | 0.630 | 0.497–0.799 |
| Live wolves only | Year-specific mass | Season-adjusted NB GAMM | −0.352 | 0.348 | 0.311 | 0.703 | 0.356–1.390 |

**Note:** The year random-effect SD in the all-wolf body-mass model was 0.59 kg. Year-specific predictions were retained as a sensitivity analysis because interannual variation could potentially provide information for distinguishing body mass from season. β represents the coefficient for standardized predicted body mass; SE = standard error; IRR = incidence rate ratio.
